# Infralimbic projections to the nucleus accumbens shell and amygdala regulate the encoding of cocaine extinction learning

**DOI:** 10.1101/2022.10.20.513048

**Authors:** Kelle E. Nett, Alexa R. Zimbelman, Matthew S. McGregor, Vanessa Alizo Vera, Molly R. Harris, Ryan T. LaLumiere

## Abstract

Prior evidence indicates that the infralimbic cortex (IL) mediates the ongoing inhibition of cocaine seeking following self-administration and extinction training in rats, specifically through projections to the nucleus accumbens (NA) shell. Our own data indicate that IL activity immediately following an unreinforced lever press is critical for encoding the extinction contingencies in such procedures. Whether extinction encoding requires activity in the IL exclusively or also activity in its outputs, such as those to the NAshell and amygdala, is unknown. To address this issue, we used a closed-loop optogenetic approach in female and male Sprague-Dawley rats to silence IL-NAshell or IL-amygdala activity following an unreinforced lever press during extinction training. Optical illumination (20 s) was given either immediately after a lever press or following a 20 s delay. IL-NAshell inhibition immediately following an unreinforced lever press increased lever pressing during extinction training and impaired retention of extinction learning, as assessed during subsequent extinction sessions without optical inhibition. Likewise, IL-amygdala inhibition given in the same manner impaired extinction retention during sessions without inhibition. Control experiments indicate that critical encoding of extinction learning does not require activity in these pathways beyond the initial 20 s post-lever press period, as delayed IL-NAshell and IL-amygdala inhibition had no effect on extinction learning. These results suggest that a larger network extending from the IL to the NAshell and amygdala is involved in encoding extinction contingencies following cocaine self-administration.

**Significance Statement:** Infralimbic cortex (IL) activity following an unreinforced lever press during extinction learning encodes the extinction of cocaine-seeking behavior. However, the larger circuitry controlling such encoding has not been investigated. Using closed-loop optogenetic pathway targeting, we found that inhibition of IL projections to the nucleus accumbens (NA) shell and to the amygdala impaired the extinction of cocaine seeking. Importantly, these effects were only observed when activity was disrupted during the first 20 s post-lever press and not when given following a 20 s delay. These findings suggest that successful cocaine extinction encoding requires activity across a larger circuit beyond simply inputs to the IL.

## Introduction

Previous findings suggest that the infralimbic cortex (IL), the ventral portion of the rodent medial prefrontal cortex, regulates the extinction and ongoing inhibition of cocaine seeking. Evidence indicates that pharmacological inhibition and activation of the IL following cocaine extinction training sessions impairs and enhances, respectively, the consolidation of extinction learning (LaLumiere et al., 2010). As extinction learning involves detection of a prediction error following an instrumental response, it was hypothesized that important encoding of this error would occur immediately following the unreinforced lever press. Consistent with this, recent work from our laboratory indicates that optogenetic IL cell body inhibition given during the 20 s immediately following an unreinforced lever press impairs the encoding of the extinction of cocaine seeking (Gutman et al., 2017). Similar inhibition given in a pseudo-random manner throughout the extinction session had no effect on extinction learning, indicating that it is not general IL activity but rather specific windows of IL activity that are necessary for normal extinction learning. However, whether such encoding depends strictly on IL activity or involves a larger circuitry including IL projections to downstream structures is unknown.

Following extinction training, IL activity is important for suppressing cue-driven cocaine seeking (Augur et al., 2016; Muller Ewald et al., 2019). However, the IL sends dense projections to the nucleus accumbens (NA) shell, a critical hub for circuits that underlie motivated behaviors such as drug seeking (Vertes, 2004; Floresco, 2015; Gibson et al., 2019). Indeed, evidence indicates chemogenetic activation of the IL-NAshell pathway attenuates cue-induced cocaine seeking after extinction training (Augur et al., 2016). Whether this pathway is necessary for extinction encoding is unknown, as the IL-NAshell pathway may simply serve as a motor output for the extinction learning that has been stored in the IL itself. Therefore, the present experiment examined whether activity in IL projections to downstream structures, such as those to the NAshell, is important for encoding the extinction of cocaine seeking.

Although studies examining IL control over the inhibition of cocaine seeking have mainly focused on NAshell outputs, the IL also sends projections to the amygdala, a region implicated in extinction behaviors for tone fear conditioning (Maren and Quirk, 2004). Anatomical studies indicate that the IL projects to the amygdala, with dense projections to GABAergic intercalated cells (ITCs) located in the medial zone between the basolateral amygdala (BLA) and central amygdala (Pinard et al., 2012). ITC activity is important for the expression of the extinction of conditioned fear, and evidence suggests that this is mediated by projections from the IL (Quirk et al., 2003; Likhtik et al., 2008; Duvarci and Pare, 2014; Bukalo et al., 2015; Bloodgood et al., 2018). Considering that the BLA itself promotes both fear expression and cocaine seeking (Kruzich and See, 2001; Maren, 2003), these findings suggest that IL inputs to this population of medial ITCs (mITCs) are involved in the extinction of cocaine seeking. However, other evidence suggests that activity in IL projections to the NAshell and BLA have different roles in the punishment-induced suppression of ethanol seeking (Halladay et al., 2019), raising the possibility that such distinctions are present in the extinction of cocaine seeking as well. Thus, whether the IL-amygdala and IL-NAshell activity are similarly important for extinction encoding or whether such encoding is specific to one pathway remains unclear.

To address these questions, the present work used a closed-loop optogenetic approach to inhibit IL terminals in the NAshell or amygdala immediately following unreinforced lever presses during early extinction training after cocaine self-administration. In contrast to prior work using only males (Gutman et al., 2017), the present study incorporated both sexes, making it possible to determine whether these pathways play qualitatively different roles between sexes in the extinction of cocaine self-administration. Overall, the results suggest that both pathways are involved in the extinction encoding for cocaine seeking, with similar results in both sexes.

## Methods

### Subjects

Female and male Sprague-Dawley rats (200-225 g and 225-250 g, respectively, at the time of first surgery; Envigo; *n* = 106) were used for this study. All rats were single-housed in a temperature-controlled environment under a 12 h light/dark cycle (lights on at 07:00) and allowed to acclimate to the vivarium at least 2 d before surgery. All procedures followed the National Institutes of Health guidelines for care of laboratory animals and were approved by the University of Iowa Institutional Animal Care and Use Committee.

### Surgery

Rats were anesthetized with 3-5% isoflurane. Meloxicam (2 mg/kg, s.c.) was administered as an analgesic before surgery as well as 24 h after surgery. Rats also received sterile saline (3 ml, s.c.) after surgery for rehydration. All rats underwent two surgeries separated by 2 weeks. Virus was injected during the first surgery, and catheters and optical fibers were implanted during the second surgery.

For catheter implantation, a 15 cm rounded tip rat jugular vein catheter (SAI Infusion Technologies) with suture beads 3.0 and 3.5 cm from the rounded tip was inserted into the right jugular vein. The opposite end of the catheter was externalized between the shoulder blades and connected to a harness with a 22-gauge guide cannula, which was used for the delivery of cocaine. Catheters were flushed 6 d per week with 0.1 ml of heparinized saline and glycerol to ensure catheter patency. Rats received antibiotics (Baytril; 2.5 mg/kg, s.c.) the day of catheter implantation and for 12 d following surgery.

For virus injection and optical fiber implantation, rats were placed in a small animal stereotax (Kopf Instruments) and injected with virus (0.3 μl; AAV5-CaMKIIα-eArchT3.0-eYFP or AAV5-CaMKIIα-eYFP) delivered bilaterally into the IL (AP +3.0, ML +0.6, DV -5.5) through double-barreled 33-gauge injectors (1.2 mm center-to-center distance; Plastics One) at a rate of 0.1 μl/min. Injectors were left in place for 7 min to allow diffusion of the virus. Rats were also implanted with indwelling optical fibers bilaterally targeting the NAshell (AP +1.2, ML +1.2, DV -7.0 at a 10° angle), mITCs between the BLA and central amygdala (AP -2.5, ML +4.9, DV -7.6 at a 5° angle), or IL (AP +3.0, ML+1.5, DV-4.5 at a 10° angle), with all angles with respect to the sagittal plane. Optical fiber implants were made in-house by gluing optical fibers (0.5 NA, Ø200 μm core; Thor Labs) into multimode stainless alloy ferrules (2.5 mm long with 230-240 μm bore; Precision Fiber Products), and the externalized end of the ferrule was polished using lapping sheets with decreasing grit (5 -0.3 μm; Thor Labs). Dust caps were maintained on the externalized end of the ferrule throughout the experiments.

### Optical illumination

During sessions in which rats received optical illumination, rats were connected to a laser (300 mW, 561 nm, OEM lasers) as previously described (Gutman et al., 2017). Laser output was measured using a power meter and adjusted to ∼10 mW at the fiber tip, based on previous work (Yizhar et al., 2011; Gutman et al., 2017).

### Cocaine self-administration

Self-administration training sessions were carried out 6 d per week in standard operant conditioning chambers, housed within sound-attenuating chambers (Med Associates) and equipped with a central reward magazine flanked by two retractable levers. Cue lights were located directly above the levers, and a 4500 Hz Sonalert module above the right lever was used as the tone generator. A house light on the opposite wall of the operant chamber was illuminated throughout the training sessions. After 24 h of food deprivation, rats were trained in an overnight session to lever press for 45 mg food pellets (Bio-serv Dustless Precision Pellets) on an FR1 schedule of reinforcement. One day after food training, rats began training 6 d per week on the 2 h cocaine self-administration task.

During cocaine self-administration, a lever press on the active (right) lever resulted in a 50 μl cocaine infusion (dissolved in 0.9% sterile saline; cocaine kindly provided by the National Institute on Drug Abuse) and the presentation of the cue light directly above the active lever and tone cues, both for 5 s. Female and male doses were 65 and 100 ug/infusion, respectively, leading to ∼0.33 mg/kg/infusion for both sexes. During the initial days of self-administration training, a timeout period (20 s) followed each infusion, during which active lever presses were recorded but had no scheduled consequence. Following at least 2 d of cocaine self-administration with ≥ 15 infusions, rats were trained on the full self-administration task, in which the active lever was retracted for 20 s (Experiments 1 and 3) or 40 s (Experiments 2 and 4) immediately following each infusion. The levers were retracted in such a manner during self-administration to familiarize the rat with lever retraction procedures that occurred during optogenetic manipulations during extinction. Self-administration completion criteria included ≥ 12 d of cocaine self-administration with ≥ 10 infusions on 10 of the days and ≥ 15 infusions on each of the final 3 d.

### Extinction

After reaching completion criteria for self-administration, rats began extinction training. Initially, rats underwent 5 d of 30 min extinction sessions in which each lever press produced lever retraction and 20 s of laser illumination, either immediately following the lever press (Experiments 1 and 3) or after a 20 s delay (Experiments 2 and 4). The levers were retracted in this manner so that rats could not press the lever during the laser illumination. Experiments 2 and 4 used a 40 s retraction so that illumination could be given in the 20-40 s window following the unrewarded lever press to determine whether activity important for encoding extended beyond the initial 20 s following an unrewarded lever press.

Following these shortened, manipulated sessions, rats underwent 7 d of 2 h extinction sessions in which each active lever press produced the lever retraction for 20 or 40 s but no laser illumination. The extinction data from these 7 d served as an index of retention of the extinction learning from the shortened sessions. The choice for 5 d of shortened extinction sessions, followed by full-length sessions, was based on previous work (LaLumiere et al., 2010; Gutman et al., 2017). This design reduces the amount of extinction learning that occurs during each shortened extinction session, thereby enabling the full-length extinction sessions to better serve as an index of retention.

### Cued reinstatement

After extinction, rats underwent cued reinstatement. To undergo reinstatement, rats needed ≥ 7 d of 2 h extinction sessions and ≤ 15 active lever presses on the 3 consecutive extinction days immediately before the reinstatement session. For cued reinstatement, active lever presses resulted in 20 s lever retraction and produced the light and tone cues previously associated with the cocaine infusion but did not produce a cocaine infusion. The specific extinction and manipulation procedures for each experiment are described below.

### Experiment 1: IL-NAshell inhibition immediately following an unreinforced lever press during extinction training

Experiment 1 examined whether post-lever press IL-NAshell activity is necessary for the extinction of cocaine seeking. Here, a viral vector expressing inhibitory opsin (eArchT) or empty vector control (eYFP) was injected into the IL and optical fibers were implanted above IL terminals in the NAshell. In this experiment, a lever press produced 20 s of lever retraction.

To confirm that increased lever pressing was not the result of any rewarding or locomotor effects of pathway illumination, we also examined whether rats would press a lever to receive inhibition of the IL-NAshell pathway. As this possibility was not examined by the original work from Gutman et al. (2017), the same experiment was also conducted for inhibition of IL cell bodies. Rats were injected with a viral vector expressing with the inhibitory opsin (eArchT) or empty vector control (eYFP) and had optical fibers implanted directly above the IL (for IL cell body inhibition) or NAshell (for IL-NAshell pathway inhibition). Rats underwent overnight food training as described above to establish lever pressing behavior. Then, rats underwent optical illumination self-administration, in which an active lever press resulted in a 20 s lever retraction, light and tone cues, and laser illumination for 20 s. Rats underwent this optical self-administration for 7 d.

### Experiment 2: Delayed IL-NAshell inhibition following an unreinforced lever press during extinction training

Experiment 2 determined whether IL-NAshell activity in the 20-40 s period following a lever press was important for extinction encoding. In this case, during self-administration, an active lever press produced a 40 s lever retraction. During the 5 d of shortened extinction sessions, active lever presses resulted in lever retraction for 40 s and laser illumination starting at 20 s post-lever press and continuing to 40 s, at which time the laser turned off and the lever was reinserted. During the following 7 d of full-length, unmanipulated extinction sessions, active lever presses only resulted in lever retraction for 40 s. Cue-induced reinstatement testing occurred as described above, with a 40 s lever retraction following each lever press.

### Experiment 3: IL-amygdala inhibition following an unreinforced lever press during extinction training

Experiment 3 examined whether IL projections to the amygdala are involved in the extinction of cocaine seeking. Except for illumination being provided to the IL terminals in the amygdala, all procedures were identical to Experiment 1.

### Experiment 4: Delayed IL-amygdala inhibition following an unreinforced lever press during extinction training

Experiment 4 investigated whether activity in the IL-amygdala pathway during the 20-40 s period after an active lever press was necessary for normal extinction encoding. Thus, all procedures were the same as Experiment 2 except that inhibition was provided to the IL-amygdala pathway.

### Histology

Rats were overdosed with sodium pentobarbital (100 mg/kg, i.p.) and transcardially perfused with 60 ml of PBS (pH 7.4), followed by 60 ml of 4% paraformaldehyde in PBS. Brains were stored in 4% paraformaldehyde for 48 h before sectioning. Brains were coronally sectioned (75 μm) and mounted on gelatin-coated slides to either be stained with Cresyl violet or viewed under a fluorescent microscope. Optical fiber termination points were visualized on Cresyl violet-stained sections under a light microscope according to the Paxinos and Watson atlas (Paxinos, 2007). Sections were viewed under a fluorescent microscope to verify viral expression. Rats with misplaced virus expression or optical probes were excluded from analysis.

### Statistical analysis

Active lever presses and infusions during the last 3 d of cocaine self-administration were analyzed using a two-way, repeated-measures ANOVA with day as the within-subjects variable and manipulation (eYFP vs eArchT) as the between-subjects variable. The same analysis was also used to analyze active lever presses during the shortened/manipulated extinction sessions, cue-induced reinstatements, and self-administration of optogenetic illumination. For cue-induced reinstatement, the extinction baseline (average active lever presses over the last 3 d of extinction) was compared to active lever pressing during the cue-induced reinstatement test for the within-subject variable. Although not fully powered by sex, each two-way, repeated-measures ANOVA was also run separately for females and males as a preliminary analysis to identify potential areas in which differences may emerge and in accordance with NIH policy on sex as a biological variable.

To analyze active lever presses during the 7 d of unmanipulated, full-length extinction sessions, non-linear mixed effects modeling with rat as a random variable was performed using the “lme4” package (v1.1-30) within R Statistical Software (v4.2.1; R Core Team 2021). This type of analysis better represents the exponential shape of the extinction curve compared to the traditional repeated-measures ANOVA (Pinheiro, 2000). However, the 5 d of shortened extinction sessions and female-only and male-only analyses were done using a two-way, repeated-measures ANOVA as non-linear mixed effects modeling requires more subjects and data points across time. In some instances, rats that successfully completed the extinction training were unable to complete reinstatement testing due to lost headcaps, illness, and death, leading to a decrease in the n in some experiments. For all repeated-measures ANOVAs, the Greenhouse-Geisser correction was used if the assumption of sphericity was violated. Unless otherwise stated, data were analyzed in GraphPad Prism 9.0.0 (GraphPad Software, La Jolla, CA).

## Results

As shown in Table 1, the analyses of the self-administration data indicate that there were no pre-existing differences between the groups. Additionally, as the pattern of effects was the same in both females and males throughout the experiments, the statistical analyses disaggregated by sex for Experiments 1, 2, 3, and 4 were grouped together in Tables 2-5, respectively.

**Table 1.** Statistics from the last 3 d of self-administration for each experiment.

| **Measure** | **Effect** | **Experiment 1** | **Experiment 2** | **Experiment 3** | **Experiment 4** |
| --- | --- | --- | --- | --- | --- |
| Active lever presses | **Manipulation** | $F_{1, 28} = 0.64,$<br>$p = 0.43$ | $F_{1, 17} = 0.16,$<br>$p = 0.69$ | $F_{1, 20} = 0.34,$<br>$p = 0.57$ | $F_{1, 14} = 1.22,$<br>$p = 0.29$ |
| | **Day** | $F_{2, 56} = 1.13,$<br>$p = 0.33$ | $F_{1.57, 26.76} = 0.87,$<br>$p = 0.41$ | $F_{1.88, 37.50} = 2.1,$<br>$p = 0.14$ | $F_{1.73, 24.18} = 0.51,$<br>$p = 0.58$ |
| | **Interaction** | $F_{2, 56} = 0.16,$<br>$p = 0.85$ | $F_{2, 34} = 0.78,$<br>$p = 0.47$ | $F_{2, 40} = 0.46,$<br>$p = 0.64$ | $F_{2, 28} = 1.01,$<br>$p = 0.38.$ |
| Infusions | **Manipulation** | $F_{1, 28} = 0.88,$<br>$p = 0.36$ | $F_{1, 17} = 0.26,$<br>$p = 0.62$ | $F_{1, 20} = 1.09,$<br>$p = 0.31$ | $F_{1, 14} = 1.03,$<br>$p = 0.33$ |
| | **Day** | $F_{1.33, 37.24} = 0.99,$<br>$p = 0.35$ | $F_{1.38, 23.51} = 0.43,$<br>$p = 0.58$ | $F_{1.98, 39.66} = 1.38,$<br>$p = 0.26$ | $F_{2, 28} = 0.56,$<br>$p = 0.58$ |
| | **Interaction** | $F_{2, 56} = 0.23,$<br>$p = 0.80$ | $F_{2, 34} = 1.12,$<br>$p = 0.34$ | $F_{2, 40} = 0.35,$<br>$p = 0.71$ | $F_{2, 28} = 1.11,$<br>$p = 0.34)$ |
| Mg/kg cocaine | **Manipulation** | $F_{1, 28} = 0.88,$<br>$p = 0.36$ | $F_{1, 17} = 0.23,$<br>$p = 0.64$ | $F_{1, 20} = 1.09,$<br>$p = 0.31$ | $F_{1, 14} = 1.60,$<br>$p = 0.23$ |
| | **Day** | $F_{1.33, 37.24} = 0.99,$<br>$p = 0.35$ | $F_{1.34, 22.81} = 0.57,$<br>$p = 0.51$ | $F_{1.98, 39.66} = 1.38,$<br>$p = 0.26$ | $F_{1.85, 25.84} = 0.40,$<br>$p = 0.66$ |
| | **Interaction** | $F_{2, 56} = 0.23,$<br>$p = 0.80$ | $F_{2, 34} = 0.92,$<br>$p = 0.41$ | $F_{2, 40} = 0.35,$<br>$p = 0.71$ | $F_{2, 28} = 1.43,$<br>$p = 0.26$ |

**Table 2.** Female and male statistics from Experiment 1: IL-NAshell, 0-20 s inhibition.

| **Behavior** | **Effect** | **Female** | **Male** |
| --- | --- | --- | --- |
|  |  | (eYFP n = 7; eArchT n = 7) | (eYFP n = 7; eArchT n = 9) |
| Manipulated (30 min) extinction | **Manipulation** | $F_{1, 12} = 16.71, p < 0.01$ | $F_{1, 14} = 3.82, p = 0.07$ |
| | **Day** | $F_{3.36, 40.26} = 6.85, p < 0.001$ | $F_{2.23, 31.28} = 2.09, p = 0.14$ |
| | **Interaction** | $F_{4, 48} = 1.55, p = 0.20$ | $F_{4, 56} = 1.02, p = 0.40$ |
| Unmanipulated (2 h) extinction | **Manipulation** | $F_{1, 12} = 3.39, p = 0.09$ | $F_{1, 14} = 4.68, p = 0.05$ |
| | **Day** | $F_{3.35, 40.19} = 12.24, p < 0.0001$ | $F_{2.45, 34.23} = 8.86, p < 0.001$ |
| | **Interaction** | $F_{6, 72} = 2.41, p = 0.04$ | $F_{6, 84} = 1.59, p = 0.16$ |
| Cued reinstatement | **Manipulation** | $F_{1, 12} = 0.70, p = 0.42$ | $F_{1, 14} = 0.58, p = 0.46$ |
| | **Day** | $F_{1, 12} = 27.36, p < 0.001$ | $F_{1, 14} = 24.2, p < 0.001$ |
| | **Interaction** | $F_{1, 12} = 0.53, p = 0.29$ | $F_{1, 14} = 0.30, p = 0.59$ |
| Pathway inhibition "self-administration" |  | (eYFP n = 2; eArchT n = 3) | (eYFP n = 2; eArchT n = 4) |
| | **Manipulation** | $F_{1, 3} = 0.01, p = 0.95$ | $F_{1, 4} = 0.06, p = 0.82$ |
| | **Day** | $F_{2.00, 5.99} = 0.87, p = 0.47$ | $F_{1.61, 6.42} = 0.233, p = 0.17$ |
| | **Interaction** | $F_{6, 18} = 1.57, p = 0.21$ | $F_{6, 24} = 0.56, p = 0.76$ |

**Table 3.** Female and male statistics from Experiment 2: IL-NAshell, delayed inhibition.

| **Behavior** | **Effect** | **Female**<br>(eYFP n = 4; eArchT n = 4) | **Male**<br>(eYFP n = 6; eArchT n = 5) |
| --- | --- | --- | --- |
| Manipulated (30 min)<br>extinction | **Manipulation** | $F_{1,6} = 0.58, p = 0.47$ | $F_{1,9} = 0.032, p = 0.86$ |
| | **Day** | $F_{2.45, 14.72} = 1.42, p = 0.28$ | $F_{2.27, 20.45} = 4.12, p = 0.03$ |
| | **Interaction** | $F_{4,24} = 1.17, p = 0.35$ | $F_{4,36} = 0.12, p = 0.97$ |
| Unmanipulated (2 h)<br>extinction | **Manipulation** | $F_{1,6} = 0.41, p = 0.54$ | $F_{1,8} = 0.57, p = 0.47$ |
| | **Day** | $F_{2.03, 12.20} = 13.65, p < 0.001$ | $F_{1.64, 13.14} = 10.42, p < 0.01$ |
| | **Interaction** | $F_{6,36} = 1.49, p = 0.21$ | $F_{6,48} = 0.27, p = 0.95$ |
| Cued reinstatement | **Manipulation** | $F_{1,6} = 0.58, p = 0.48$ | $F_{1,8} = 1.69, p = 0.23$ |
| | **Day** | $F_{1,6} = 25.31, p < 0.01$ | $F_{1,8} = 25.21, p < 0.01$ |
| | **Interaction** | $F_{1,6} = 1.19, p = 0.32$ | $F_{1,8} = 2.60, p = 0.15$ |

**Table 4.** Female and male statistics from Experiment 3: IL-amygdala, 0-20 s inhibition.

| **Behavior** | **Effect** | **Female** | **Male** |
| --- | --- | --- | --- |
|  |  | (eYFP n = 6; eArchT n = 6) | (eYFP n = 4; eArchT n = 6) |
| Manipulated (30 min)<br>extinction | **Manipulation** | $F_{1,10} = 2.59, p = 0.14$ | $F_{1,8} = 0.43, p = 0.53$ |
| | **Day** | $F_{2.78, 27.75} = 12.05, p < 0.0001$ | $F_{1.71, 13.69} = 4.08, p = 0.05$ |
| | **Interaction** | $F_{4,40} = 0.42, p = 0.80$ | $F_{4,32} = 0.19, p = 0.94$ |
| Unmanipulated (2 h)<br>extinction | **Manipulation** | $F_{1,10} = 8.34, p = 0.02$ | $F_{1,8} = 3.68, p = 0.09$ |
| | **Day** | $F_{3.66, 36.57} = 3.27, p = 0.02$ | $F_{2.54, 20.33} = 5.14, p = 0.01$ |
| | **Interaction** | $F_{6,60} = 0.92, p = 0.49$ | $F_{6,48} = 1.92, p = 0.10$ |
| Cued reinstatement | **Manipulation** | $F_{1,10} = 3.04, p = 0.11$ | $F_{1,8} = 0.10, p = 0.76$ |
| | **Day** | $F_{1,10} = 32.63, p = 0.0002$ | $F_{1,8} = 21.23, p = 0.002$ |
| | **Interaction** | $F_{1,10} = 1.17, p = 0.31$ | $F_{1,8} = 0.04, p = 0.85$ |

**Table 5.** Female and male statistics from Experiment 4: IL-amygdala, delayed inhibition.

| **Behavior** | **Effect** | **Female** | **Male** |
| --- | --- | --- | --- |
|  |  | (eYFP n = 4; eArchT n = 3) | (eYFP n = 4; eArchT n = 5) |
| Manipulated (30 min)<br>extinction | **Manipulation** | $F_{1,5} = 1.45, p = 0.28$ | $F_{1,7} = 0.002, p = 0.96$ |
| | **Day** | $F_{2.30, 11.52} = 24.72, p < 0.0001$ | $F_{2.68, 18.78} = 11.60, p < 0.001$ |
| | **Interaction** | $F_{4,20} = 0.82, p = 0.53$ | $F_{4,28} = 1.15, p = 0.35$ |
| Unmanipulated (2 h)<br>extinction | **Manipulation** | $F_{1,5} = 0.06, p = 0.82$ | $F_{1,7} = 0.12, p = 0.74$ |
| | **Day** | $F_{3.22, 16.10} = 7.17, p = 0.003$ | $F_{2.57, 17.95} = 7.23, p = 0.003$ |
| | **Interaction** | $F_{6,30} = 0.71, p = 0.64$ | $F_{6,42} = 1.55, p = 0.19$ |
| Cued reinstatement | **Manipulation** | $F_{1,5} = 0.10, p = 0.40$ | $F_{1,7} = 0.70, p = 0.43$ |
| | **Day** | $F_{1,5} = 6.95, p = 0.05$ | $F_{1,7} = 51.36, p < 0.001$ |
| | **Interaction** | $F_{1,5} = 0.10, p = 0.76$ | $F_{1,7} = 0.79, p = 0.40$ |

### Experiment 1

In Experiment 1, IL-NAshell inhibition was given for 20 s immediately following an unreinforced lever press during the 5 d of shortened extinction (Figure 1A-C). Figure 1D shows active lever presses across extinction sessions. Analysis of active lever presses during the shortened, manipulated extinction sessions revealed main effects of inhibition and day and a trend toward an interaction (*F*_1, 28_ = 3.08, *p* < 0.01; *F*_2.82, 78.85_ = 7.91, *p* < 0.001; *F*_4, 122_ = 2.20, *p* = 0.07, respectively). Thus, immediate post-lever press inhibition of IL-NAshell increased active lever pressing during sessions in which manipulations were given. Inhibition of IL-NAshell also impaired extinction retention, as assessed during the full-length, unmanipulated extinction sessions using non-linear mixed effect modeling with rat as a random effect. Analysis of active lever presses revealed main effects of inhibition, extinction day, and extinction rate (*t*_86.82_ = -4.69, *p* < 0.0001; *t*_176_ = -7.98, *p* < 0.001; *t*_176_ = 5.53, *p* < 0.0001, respectively). Although there was a significant interaction between day and inhibition, there was not a significant interaction between manipulation and the extinction rate (*t*_176_ = 2.24, *p* = 0.03; *t*_176_ = -1.09, *p* = 0.28, respectively). Both groups sufficiently extinguished lever pressing, and there were no differences in cue-induced reinstatement of cocaine seeking (Figure 1E). Both groups reinstated active lever pressing to the drug-associated cues (*F*_1, 28_ = 51.91, *p* < 0.0001), but there was no main effect of inhibition and no interaction (*F*_1, 28_ = 1.15, *p* = 0.29; *F*_1, 28_ = 0.69, *p* = 0.41, respectively).

**Figure 1.**
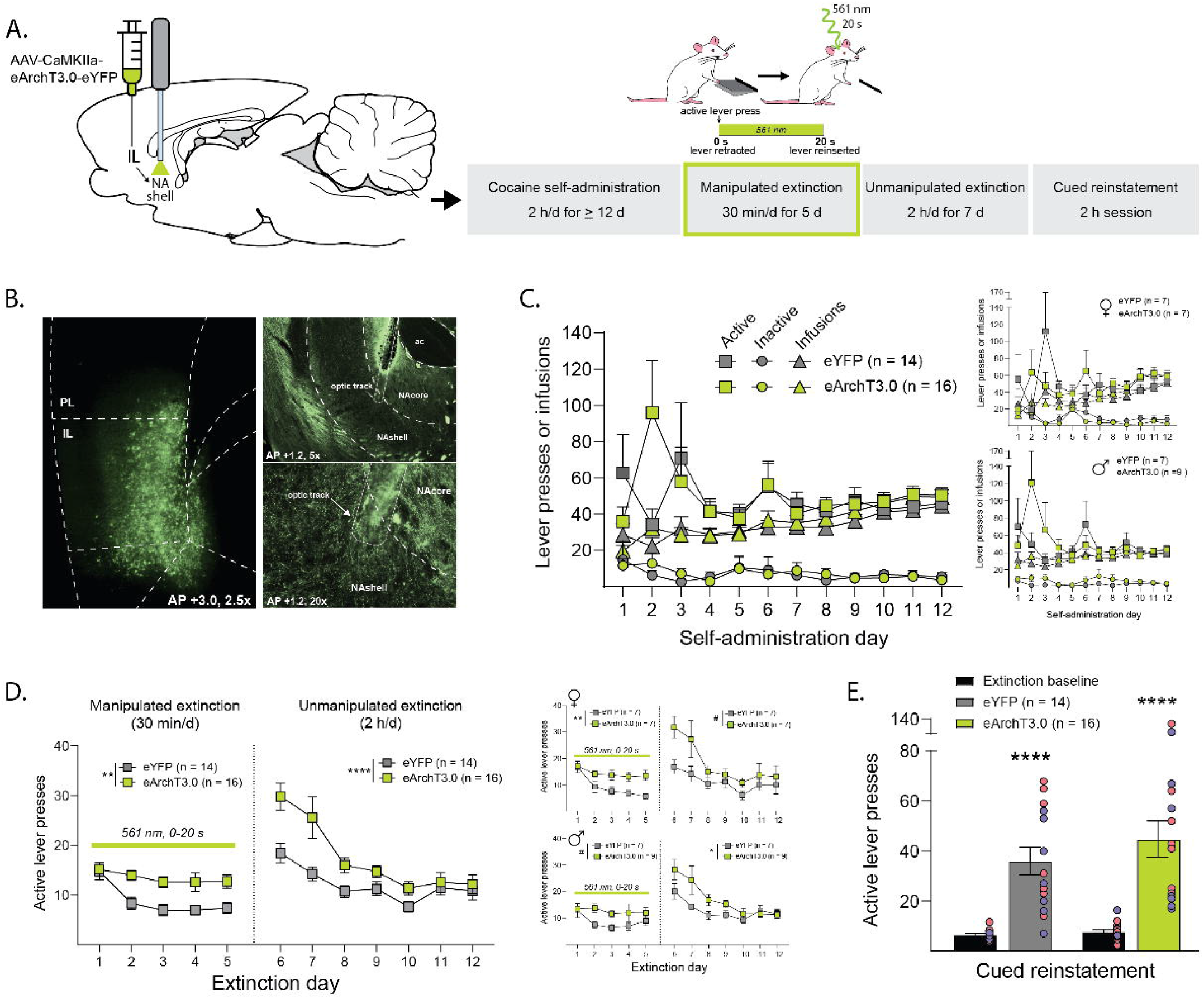
Impaired cocaine extinction learning with immediate post-lever IL-NAshell inhibition. **A**. *Left*, A viral vector containing the inhibitory opsin, eArchT3.0, was injected upstream into the IL and optical fibers were implanted into the NAshell to target IL terminals. *Right*, after recovery from surgery, rats underwent daily 2 h cocaine self-administration, followed by 5 d of 30 min manipulated extinction sessions, in which each active lever press resulted in 20 s lever retraction and laser illumination for the duration of the lever retraction. Rats then underwent 7 d of full-length (2 h) unmanipulated extinction sessions to assess the retention of the extinction learning, followed by cued reinstatement. **B**. Representative fluorescent images depicting (*left*) virus expression in IL cell bodies and (*right*) virus expression in IL terminals in the NAshell where the optical fiber terminates. **C**. Active and inactive lever presses and infusions during cocaine self-administration did not differ between groups and were similar in female (*right, top*) and male (*right, bottom*) rats. **D**. Immediate post-lever press inhibition of the IL-NAshell pathway increased active lever presses during manipulated sessions and unmanipulated sessions, though it did not impair the ability of these rats to eventually extinguish lever pressing. A similar effect was observed in female (*right, top*) and male (*right, bottom*) rats. **E**. Both groups had increased lever pressing during cued reinstatement with no differences between eArchT3.0 and eYFP rats. Individual data points for female and male rats are depicted in red and blue circles, respectively. # *p* < 0.1, * *p* < 0.05, ** *p* < 0.01, *** *p* < 0.001, **** *p* < 0.0001

To determine whether IL inhibition or IL-NAshell inhibition alone could be responsible for the observed effects as well as those from our previous work (Gutman et al., 2017), separate groups of rats underwent 7 d of optogenetic self-administeration in which an active lever press resulted in a 20 s lever retraction and laser illumination of IL cell bodies (Figure 2A-B) or IL-NAshell pathway (Figure 2C-D). Analysis of active lever presses for IL cell body illumination across the 7 d revealed no effect of inhibition, day, or interaction (*F*_1, 6_ = 0.73, *p* = 0.43; *F*_1.79, 10.76_ = 1.81, *p* = 0.21; *F*_6, 36_ = 1.22, *p* = 0.32, respectively). Analysis of active lever presses for IL-NAshell illumination revealed a main effect of day, but no effect of inhibition and no interaction (*F*_6, 54_ = 3.30, *p* = 0.008; *F*_1, 9_ = 0.05, *p* = 0.83; *F*_6, 54_ = 1.31, *p* = 0.27, respectively). Thus, the increase in active lever pressing observed following post-lever press inhibition of IL-NAshell during the extinction of cocaine seeking does not appear to be the result of such inhibition being rewarding or enhancing locomotor activity.

**Figure 2.**
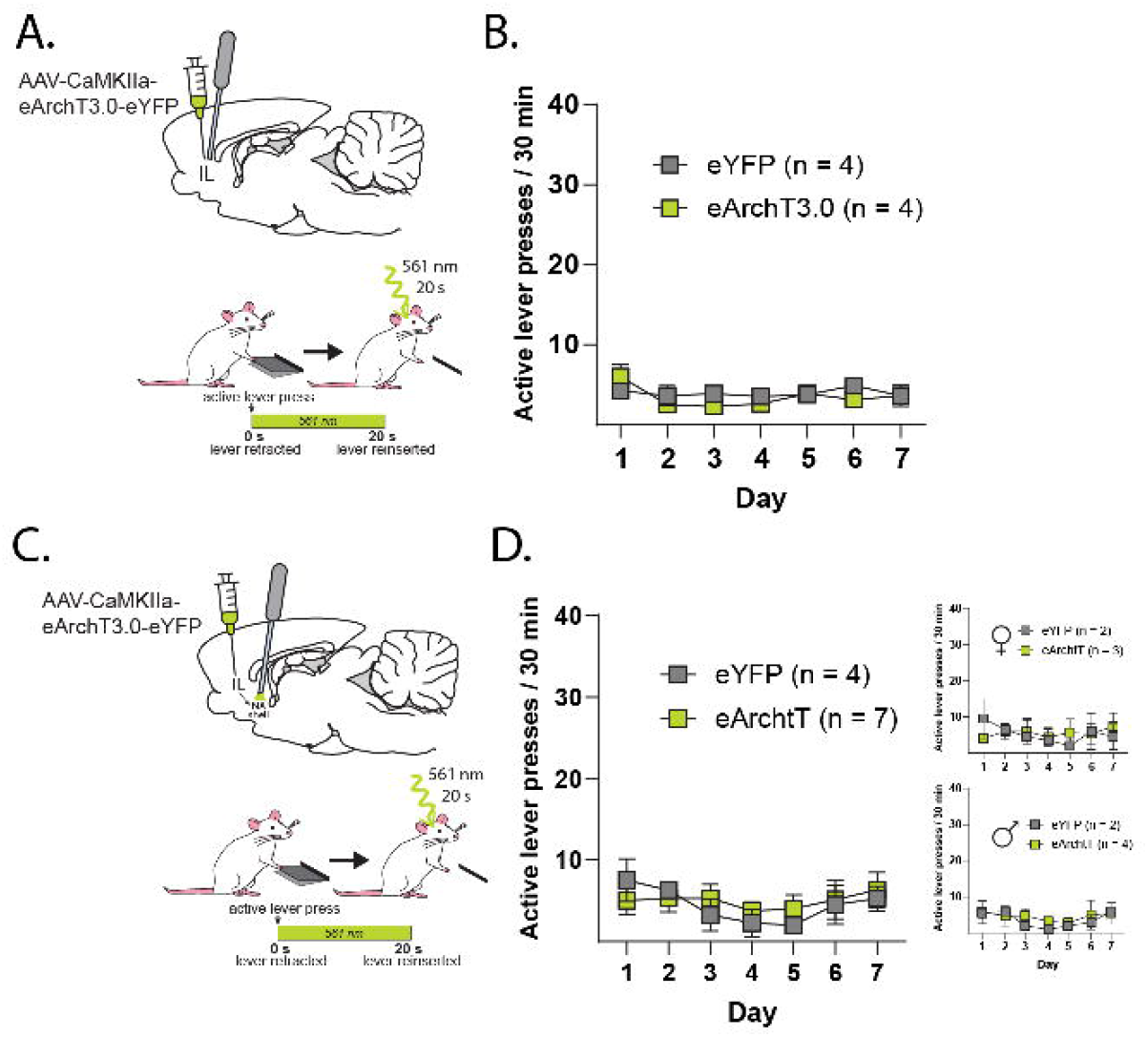
No self-administration of optogenetic inhibition of IL cell bodies or IL-NAshell projections. **A**. Male rats received intra-IL injections of a viral vector containing the inhibitory opsin eArchT or eYFP control and had optical fibers implanted above the IL. Rats were trained to press a lever for food and then began “self-administration” in which an active lever press produced a 20 s lever retraction paired with 20 s of 561 nm laser illumination. **B**. Rats did not increase lever pressing as a result of such illumination. **C**. Female and male rats received intra-IL injections of a viral vector containing the inhibitory opsin eArchT or eYFP and had optical fibers implanted directly above the NAshell. Rats were trained to press a lever for food, then began “self-administration” in which an active lever press produced a 20 s lever retraction paired with 20 s of 561 nm laser illumination. **D**. Rats did not increase lever pressing as a result of such illumination.

### Experiment 2

In Experiment 2, IL-NAshell inhibition was given during the 20-40 s period following an unreinforced lever press during the 5 d of shortened extinction (Figure 3A-C). Figure 3D shows active lever presses across extinction. Analysis of active lever pressing during the shortened extinction sessions revealed a main effect of day, but no effect of inhibition and no interaction (*F*_4, 68_ = 5.57, *p* < 0.001; *F*_1, 17_ = 0.19, *p* = 0.67; *F*_4, 68_ = 0.67, *p* = 0.62, respectively). Thus, delayed post-lever press IL-NAshell inhibition did not alter active lever pressing during extinction sessions in which post-lever press manipulations were given. Such inhibition also did not increase active lever pressing during subsequent full-length extinction sessions in which no manipulation was given, as non-linear mixed effect modeling with rat as a random effect revealed no main effect of inhibition, no interaction between extinction day and inhibition, and no interaction between inhibition and extinction rate (*t*_17.00_ = 1.21, *p* = 0.24; *t*_17.12_ = -0.49, *p* = 0.63; *t*_17.90_ = 0.21, *p* = 0.84, respectively). There were significant main effects of extinction day and extinction rate (*t*_17.12_ = -5.15, *p* < 0.0001; *t*_17.90_ = 4.18, *p* < 0.001, respectively), reflecting extinction learning in both groups. No lasting effects of delayed post-lever press inhibition during extinction were observed, as both groups successfully extinguished cocaine seeking and did not differ in cue-induced reinstatement of cocaine seeking (Figure 3E). Both groups reinstated active lever pressing to the drug-associated cues (*F*_1, 16_ = 41.56, *p* < 0.0001), with no main effect of prior inhibition and no interaction (*F*_1, 16_ = 0.36, *p* = 0.56, *F*_1, 16_ = 0.39, *p* = 0.54, respectively).

**Figure 3.**
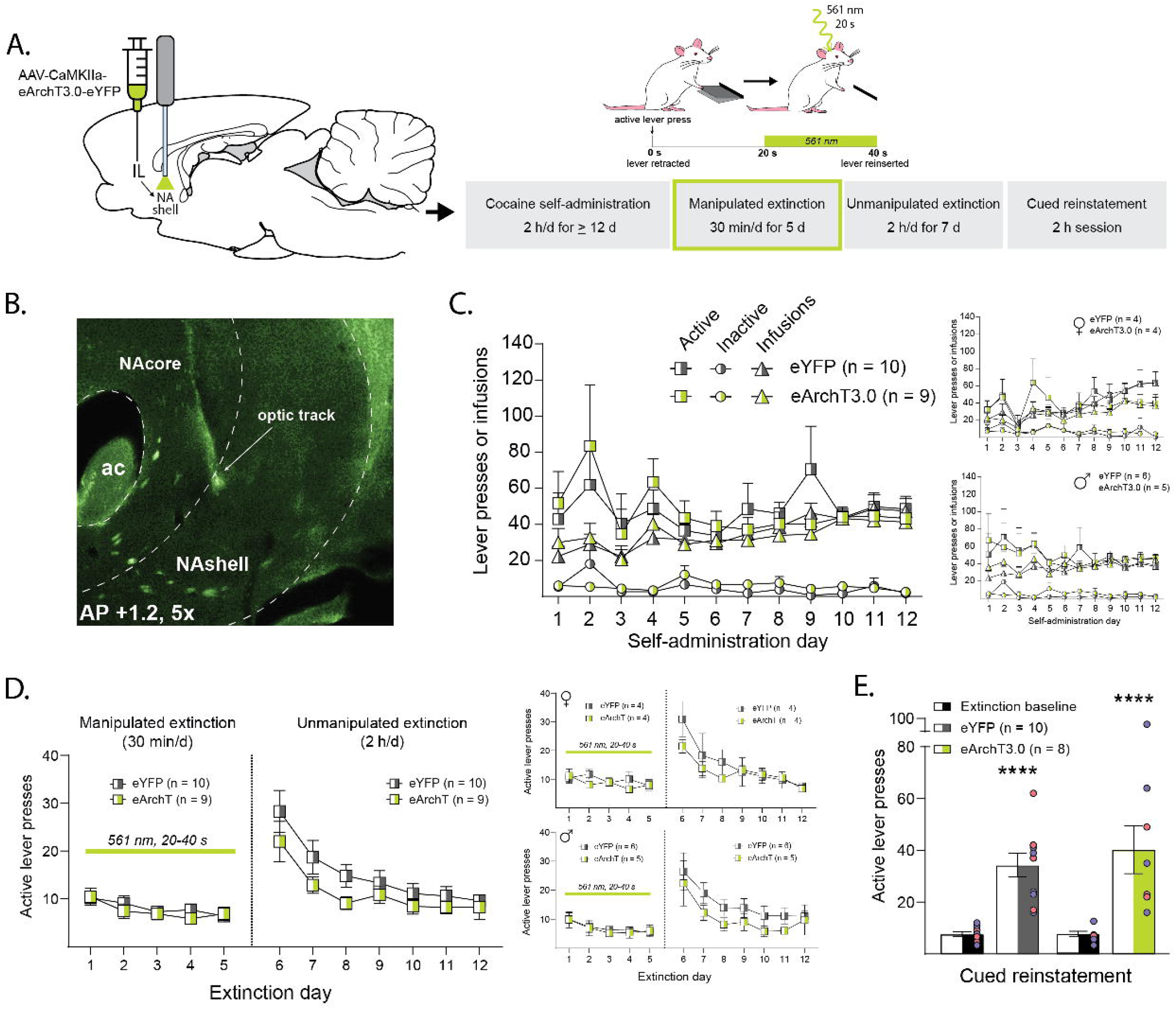
No effect of delayed post-lever press IL-NAshell inhibition on cocaine extinction learning. **A**. *Left*, A viral vector containing the inhibitory opsin eArchT3.0 (or eYFP) was injected into the IL and optical fibers were implanted into the NAshell to target IL terminals. *Right*, After recovery from surgery, rats underwent daily 2 h cocaine self-administration, followed by 5 d of 30 min manipulated extinction sessions, in which each lever press resulted in 40 s lever retraction and laser illumination starting 20 s after a lever press/lever retraction and continuing until the lever was reinserted at 40 s. Rats then underwent 7 d of full-length (2 h) unmanipulated extinction sessions to assess the retention of the extinction learning, followed by cued reinstatement. **B**. Representative fluorescent image depicting virus expression in IL terminals in the NAshell in which the optical fiber terminates. **C**. Active and inactive lever presses and infusions during cocaine self-administration did not differ between groups and were similar in female (*right, top*) and male (*right, bottom*) rats. **D**. Delayed post-lever press inhibition of IL-NAshell had no effect on active lever presses during manipulated sessions and unmanipulated sessions. Similar results were observed in female (*right, top*) and male (*right, bottom*) rats. **E**. Both groups had increased lever pressing during cued reinstatement with no differences between eArchT3.0 and eYFP rats. Individual data points for female and male rats are depicted in red and blue circles, respectively. # *p* < 0.1, * *p* < 0.05, ** *p* < 0.01, *** *p* < 0.001, **** *p* < 0.0001

### Experiment 3

Experiment 3 examined whether IL-amygdala inhibition given for 20 s immediately following an unreinforced lever press impaired extinction (Figure 4A-C). Figure 4D shows active lever presses across extinction sessions. Analysis of active lever presses during the shortened extinction sessions with inhibition revealed a trend toward a main effect of inhibition, a main effect of day, and no significant interaction (*F*_1, 20_ = 3.02, *p* = 0.10; *F*_2.33, 46.54_ = 15.94, *p* < 0.0001; *F*_4, 80_ = 0.27, *p* = 0.90, respectively). Thus, although post-lever press inhibition of IL-amygdala increased active lever presses during the shortened extinction sessions, this increase did not reach statistical significance.

**Figure 4.**
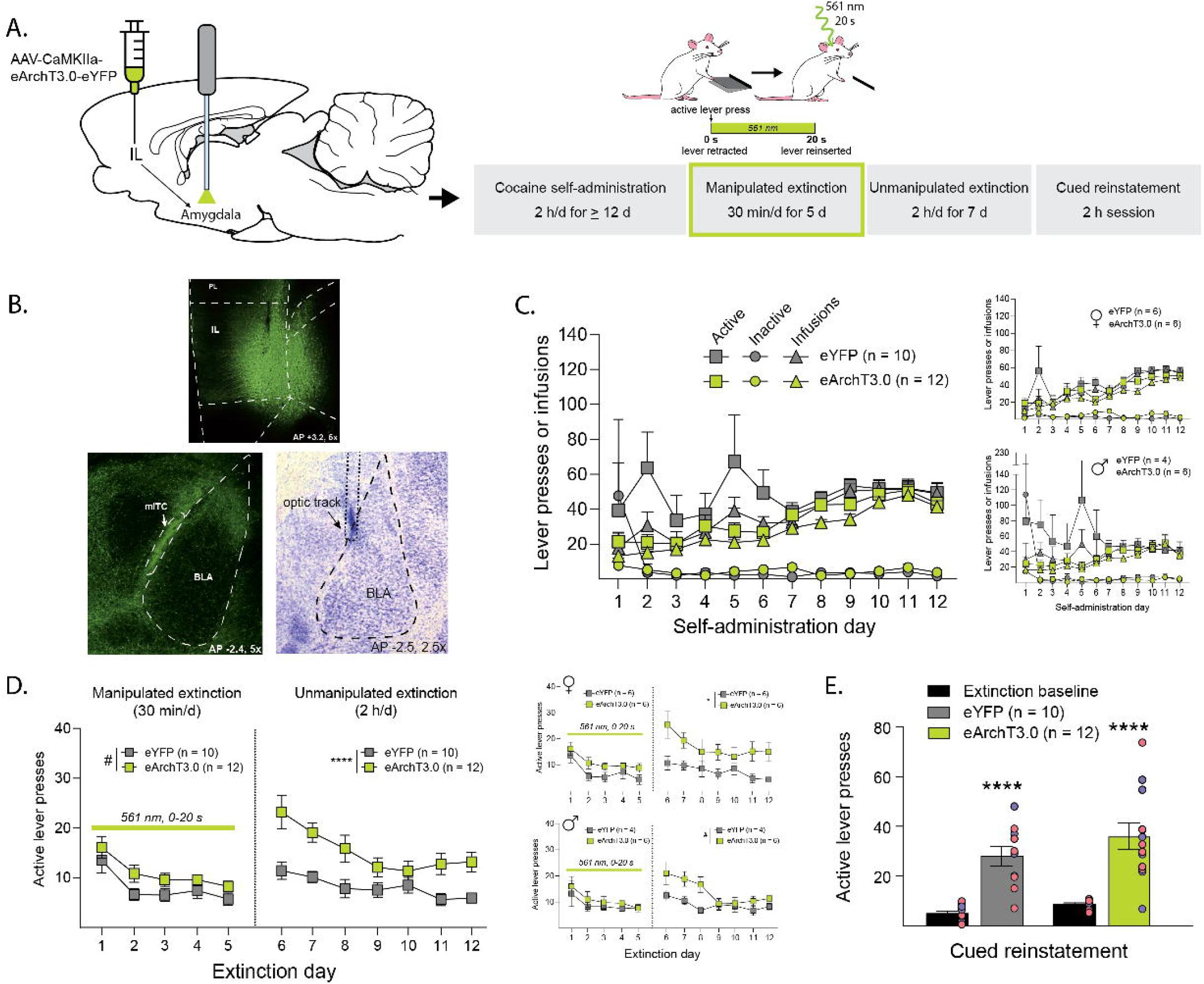
Impaired retention of cocaine extinction learning with immediate post-lever press IL-amygdala inhibition. **A**. *Left*, A viral vector containing the inhibitory opsin, eArchT3.0 (or eYFP), was injected into the IL and optical fibers were implanted above mITCs, between the BLA and central amygdala, to target IL terminals. *Right*, after recovery from surgery, rats underwent daily 2 h cocaine self-administration, followed by 5 d of 30 min manipulated extinction sessions, in which each active lever press resulted in 20 s lever retraction and laser illumination for the duration of the lever retraction. Rats then underwent 7d of full-length (2 h) unmanipulated extinction sessions to assess extinction learning retention, followed by cued reinstatement. **B**. Representative fluorescent image depicting (*top*) virus expression in IL cell bodies, (*bottom, left*) virus expression of IL terminals in the amygdala, and (*bottom, right*) the optical fiber targeting mITCs. **C**. Active and inactive lever presses and cocaine infusions during self-administration did not differ between groups and were similar in female (*right, top*) and male (*right, bottom*) rats. **D**. Immediate post-lever press inhibition of the IL-amygdala pathway had no effect on active lever presses during manipulated sessions but increased active lever presses during unmanipulated sessions. A similar effect was observed in female (*right, top*) and male (*right, bottom*) rats. **E**. Both groups had increased lever pressing during cued reinstatement with no differences between eArchT3.0 and eYFP rats. Individual data points for female and male rats are depicted in red and blue circles, respectively. # *p* < 0.1, * *p* < 0.05, ** *p* < 0.01, *** *p* < 0.001, **** *p* < 0.0001

The subsequent full-length extinction sessions without inhibition, however, revealed impaired retention in those rats that had previously received IL-amygdala inhibition. Analysis of active lever presses during the full-length extinction sessions using non-linear mixed effects modeling with rat as a random variable revealed a main effect of day, inhibition, and extinction rate (*t*_128_ = -4.70, *p* < 0.0001; *t*_51.40_ = -4.61, *p* < 0.0001; *t*_128_ = 3.05, *p* = 0.003, respectively). There was also an interaction between day and inhibition and between inhibition and extinction rate (*t*_128_ = 2.73, *p* = 0.007; *t*_128_ = -2.27, *p* = 0.03, respectively). Both groups successfully extinguished cocaine seeking and showed similar cue-induced reinstatement of cocaine seeking (Figure 4E). Both groups reinstated active lever pressing to the drug-associated cues *(F*_1, 20_ = 55.94, *p* < 0.0001), but there was no main effect of inhibition, and no interaction (*F*_1, 20_ = 2.59, *p* = 0.12; *F*_1, 20_ = 0.41, *p* = 0.53, respectively).

### Experiment 4

In Experiment 4, a separate group of rats underwent the same procedures as Experiment 3, except that lever retraction occurred for 40 s and post-lever press inhibition during the shortened extinction sessions occurred during the 20-40 s period after an active lever press (Figure 5A-C). Figure 5D shows active lever presses across extinction sessions. Analysis of active lever presses during the shortened extinction sessions with inhibition revealed a main effect of day, no main effect of inhibition, and no significant interaction (*F*_2.90, 40.62_ = 26.24, *p* < 0.0001; *F*_1, 14_ = 0.23, *p* = 0.63; *F*_4, 56_ = 0.37, *p* = 0.83, respectively). Thus, delayed post-lever press inhibition of IL-amygdala did not alter active lever presses during the shortened extinction sessions during which inhibition was given. There was also no change in active lever pressing between groups during the subsequent full-length, extinction sessions without inhibition. Analysis of active lever pressing during the full-length extinction sessions using non-linear mixed effects modeling with rat as the random variable revealed a main effect of extinction day and extinction rate (*t*_92_ = -7.24, *p* < 0.0001; *t*_92_ = 5.91, *p* < 0.0001, respectively), reflecting extinction learning. There was no main effect of inhibition, no interaction between extinction day and inhibition, and a trend toward an interaction between inhibition and extinction rate (*t*_32.67_ = 0.37, *p* = 0.72; *t*_92_ = -1.54, *p* = 0.13; *t*_92_ = 1.84, *p* = 0.07, respectively). There were no lasting effects of delayed post-lever press IL-amygdala inhibition as both groups extinguished successfully and showed similar cue-induced reinstatement of cocaine seeking (Figure 5E). Analysis of active lever presses for cue-induced reinstatement revealed that both groups reinstated active lever pressing to the drug-associated cues (*F*_1, 14_ = 38.77, *p* < 0.0001), but there was no main effect of inhibition and no interaction (*F*_1, 14_ = 0.14, *p* = 0.72; *F*_1, 14_ = 0.03, *p* = 0.86, respectively).

**Figure 5.**
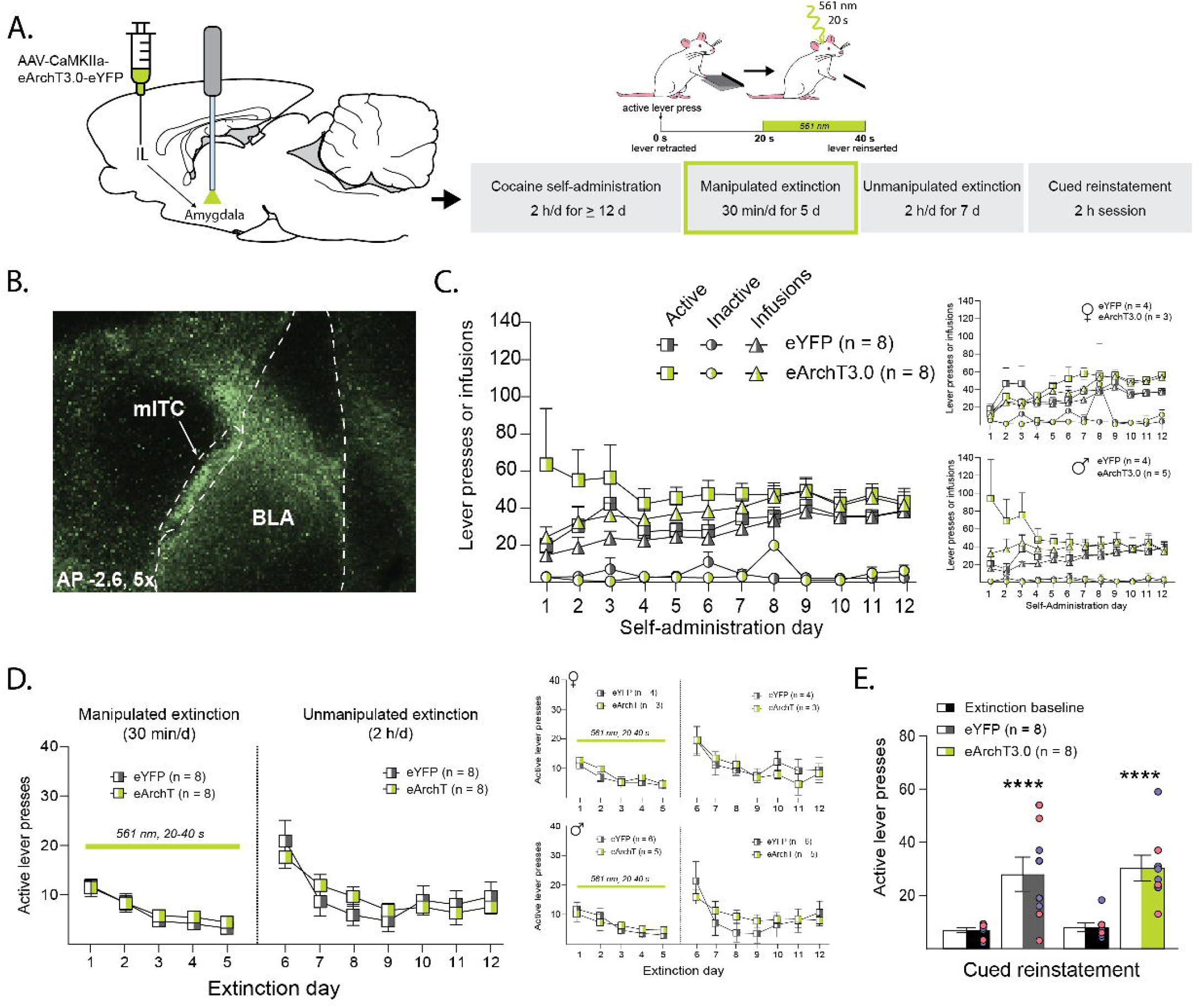
No effect of delayed post-lever IL-amygdala inhibition on cocaine extinction learning. **A**. *Left*, A viral vector containing the inhibitory opsin eArchT3.0 (or eYFP) was injected into the IL and optical fibers were implanted above mITCs, between the BLA and central amygdala, to target IL terminals. *Right*, After recovery from surgery, rats underwent daily 2 h cocaine self-administration, followed by 5 d of 30 min manipulated extinction sessions, in which each lever press resulted in 40 s lever retraction and laser illumination starting 20 s after a lever press/lever retraction and continuing until the lever was reinserted at 40 s. Rats then underwent 7 d of full-length (2 h) unmanipulated extinction sessions to assess extinction learning retention, followed by cued reinstatement. **B**. Representative fluorescent image depicting virus expression in IL terminals in the amygdala. **C**. Active and inactive lever presses during cocaine self-administration did not differ between groups and were similar in female (*right, top*) and male (*right, bottom*) rats. **D**. Delayed post-lever press inhibition of IL-amygdala had no effect on active lever presses during manipulated sessions and unmanipulated sessions. Similar results were observed in female (*right, top*) and male (*right, bottom*) rats. **E**. Both groups had increased lever pressing during cued reinstatement with no differences between eArchT3.0 and eYFP rats. Individual data points for female and male rats are depicted in red and blue circles, respectively. # *p* < 0.1, * *p* < 0.05, ** *p* < 0.01, *** *p* < 0.001, **** *p* < 0.0001

## Discussion

The present work indicates that post-lever press optogenetic inhibition of IL-NAshell and IL-amygdala projections during extinction training impaired the extinction of cocaine seeking. Specifically, post-lever press IL-NAshell inhibition resulted in increased lever pressing during sessions in which optical inhibition occurred and during subsequent unmanipulated sessions, suggesting IL projections to the NAshell play a role in early extinction learning and the retention of such learning. Post-lever press IL-amygdala inhibition produced a non-significant increase in lever pressing during sessions in which optical manipulations occurred and impaired retention of extinction learning during subsequent unmanipulated sessions. In both cases, such differences were not observed when inhibition was given in the 20-40 s period after a lever press, indicating that critical encoding does not extend beyond the initial 20 s period. Together, the present findings point to a critical window of extinction encoding for cocaine-seeking behavior that requires activity in IL projections to both downstream regions.

### Encoding of cocaine extinction contingencies during the 20 s post-lever press period

Previous work from our laboratory identified a 20 s post-lever press window during extinction learning for cocaine seeking in which IL activity is involved in encoding extinction contingencies (Gutman et al., 2017). Because extinction learning involves a prediction error following an unreinforced lever press, these findings suggest that IL activity is important for encoding the information for the extinction-based prediction error. One possible component of this prediction error is the absence of the expected rise in dopamine that occurs with a cocaine infusion. Indeed, Gutman et al. chose 20 s of optical inhibition based on evidence that dopamine levels in the brain following intravenous cocaine infusions peak within 10-20 s (Aragona et al., 2008), suggesting that activity related to the detection of the absence of the cocaine infusion would occur in a similar timeframe. However, dopamine concentrations in the NAshell following intravenous cocaine infusions remain elevated up to 90 s (Aragona et al., 2008), raising the possibility that the absence of such reinforcers may be detected, and therefore encoded, beyond the initial 20 s. The present work, thus, specifically identifies the *immediate* 20 s window after a lever press as critical for extinction encoding, as delayed inhibition of IL-NAshell and IL-amygdala pathways did not alter cocaine extinction learning.

The current findings raise an important question concerning what precisely is encoded during this post-lever press window. During extinction sessions, the previously learned instrumental contingencies are altered, such that a lever press produces no consequence (e.g., no cocaine infusion or light/tone cues). Neural signaling involved in encoding extinction learning presumably reflects the *absence* of previously expected outcomes. Such signaling may reflect the absence of the 5 s drug-associated stimuli, the absence of the intravenous cocaine infusion, or the absence of both the drug-associated stimuli and the cocaine infusion. Prior work indicates that 5 s of optogenetic IL inhibition given post-response during cued extinction of nicotine seeking (i.e., a lever press during extinction produces cues, but no nicotine infusion) has no effect on the extinction of nicotine seeking (Struik et al., 2019). Those findings suggest three possibilities: the IL is not involved in the extinction of *nicotine* seeking, the IL does not encode the change in contingency of previously drug-associated cues, or that IL activity for such encoding occurs beyond the 5 s of inhibition given. Thus, critical IL-based signaling during extinction learning may center on the absence of the drug infusion and its physiological effects, including the rise in central dopamine. Evidence suggests that the lateral habenula detects the absence of an expected reward or reward-predictive stimuli and inhibits dopamine neurons via projections to GABAergic rostromedial tegmental nucleus neurons that subsequently inhibit ventral tegmental area dopamine neurons (Jhou et al., 2009; Baker et al., 2016; Sosa et al., 2021). However, whether such signaling interacts with the extinction-encoding activity of the IL is unknown.

### IL-NAshell and IL-amygdala projections similarly regulate cocaine extinction encoding

The current work indicates that post-lever press IL-NAshell inhibition impaired extinction learning during sessions in which inhibition was given and extinction retention assessed during subsequent unmanipulated sessions. Prior studies suggest that extinction training recruits IL-NAshell projections to inhibit cocaine seeking (Augur et al., 2016; Muller Ewald et al., 2019), and accumulating evidence supports a general role for IL-NAshell signaling in updating and encoding contingencies between cues and behaviors (Nett and LaLumiere, 2021). The present findings corroborate this role, particularly as inhibition that occurred outside the initial 20 s post-lever press window did not alter extinction learning.

The present study also identified a novel role for IL-amygdala signaling in the extinction of cocaine-seeking behaviors, as optical IL-amygdala inhibition after a lever press impaired the retention of extinction learning assessed in subsequent unmanipulated sessions. Previous work found a critical role for IL-amygdala projections in the encoding of tone fear extinction (Bloodgood et al., 2018; Bukalo et al., 2021). Studies indicate a complex circuitry underlying fear extinction, in which the IL projects to ITCs, which then provide feedforward inhibition of central amygdala output neurons to decrease freezing (Peters et al., 2009; Bouton et al., 2021). Whether similar feedforward inhibition via IL projections to mITCs influences the extinction of cocaine seeking is difficult to determine, as light diffusion from the optical fiber in the present study likely illuminated IL terminals in the BLA as well as in the mITC. The mITCs also provide inputs to the BLA itself (Asede et al., 2015; Hagihara et al., 2021). Thus, whether the observed effects were due to influences on BLA activity, central amygdala activity, or both is unclear.

Nonetheless, the amygdala likely influences the extinction of cocaine seeking through a larger circuit. Within the mITCs, there are subdivisions that have functionally opposing roles depending upon whether they project onto BLA neurons that project back to the PL or IL (Hagihara et al., 2021). Moreover, the BLA itself also directly innervates the NAshell (Groenewegen et al., 1999), providing multiple paths out of the amygdala that may be involved in extinction encoding. Evidence suggests BLA-NAshell projections regulate reward-related behaviors triggered by reward-related cues, as blocking BLA-NAshell signaling impairs the ability of cocaine-related cues to reinstate cocaine-seeking behaviors (Setlow et al., 2002; Di Ciano and Everitt, 2004). Additionally, anatomical studies indicate that the IL and BLA provide converging inputs onto the same populations of NAshell neurons (Groenewegen et al., 1999; French and Totterdell, 2002, 2003), suggesting that information from both the IL and BLA are integrated within the NAshell to drive behaviors. Thus, important extinction-related encoding and even plasticity may be distributed throughout an IL-BLA-NAshell network. Future work probing amygdala involvement in the extinction of cocaine seeking through its downstream projections will help to further elucidate the intricacies of this encoding circuitry.

Of note, IL-amygdala inhibition during shortened, manipulated sessions produced a non-significant increase in active lever presses during those sessions. One possible explanation for this discrepancy with the IL-NAshell results is that the two pathways have different roles in extinction encoding. However, it is difficult to interpret the findings in that manner considering the impaired retention observed in the 7 d of full-length extinction. Indeed, this pattern (e.g., no significant effect during shortened extinction sessions but impaired retention observed on full-length extinction sessions) has been previously observed in studies using a similar methodology (LaLumiere et al., 2010). Moreover, the opposite pattern has also been observed (Gutman et al., 2017), though it is likely that those data were underpowered as the retention effects were in the expected direction. Thus, the present findings with the discrepancy during the shortened sessions likely reflect the behavioral variability that occurs within shorter behavioral sessions rather than distinct functions between the pathways.

### Lack of sex differences

Although there continues to be ongoing debate about the nature of sex differences in cocaine self-administration, the present work found similar results from IL-NAshell and IL-amygdala inhibition in females and males, as well as no sex differences in cocaine self-administration measures. That the pathway inhibition produced similar effects in females and males suggests that the underlying circuitry important for encoding cocaine extinction learning is the same between sexes. Thus, it is likely that the systems probed in the present study reflect basic learning and encoding mechanisms that are conserved between sexes.

## Conclusion

The present findings expand our understanding of IL regulation of the extinction of cocaine seeking, indicating the importance of temporally precise signaling to the NAshell and amygdala. In both cases, inhibition of the IL projections to these regions impaired the extinction encoding for cocaine seeking, and the importance of the pathway activity in both cases was limited to the 20 s immediately following an active lever press. The current results raise important questions regarding the larger circuits involved in the extinction of cocaine seeking and point to a more distributed system than previously considered.

